# Seabird calls are shaped by prosody, efficiency, and rhythmic encoding

**DOI:** 10.64898/2026.03.24.713940

**Authors:** Anna N Osiecka, Katarzyna Wojczulanis-Jakubas, Lara S Burchardt

**Affiliations:** Institute for Theoretical Biology, Humboldt-Universität zu Berlin, Germany; ENES Bioacoustics Research Laboratory, University of Saint-Etienne, France; Department of Vertebrate Ecology and Zoology, Faculty of Biology, University of Gdańsk, Poland

**Keywords:** Alle alle, brevity laws, declination, information compression, little auk, prolongation, rallentando, integer ratios

## Abstract

In the search for universals shaping acoustic communication across species, we increasingly look for patterns known from human languages and music in non-human animals. These parallels are often explored separately and with limited ecological context. Here, we take a deep dive into the temporal structure of a complex call used by the little auk (*Alle alle*), a pelagic seabird with elaborate vocal behaviour and socially complex colonial life. Based on syllable durations, intervals and silences, we examine its conformance to linguistic laws, rhythmic structure and information content. This reveals intricate problems of temporal organisation: while the calls conform not only to linguistic laws of brevity but also to the initial and final lengthening known from human prosody, these effects interact with the internal structure of the call and information carried within it. To our knowledge, this is the first time that conformance to multiple linguistic laws, exceeding simple vocal efficiency, has been described for a non-human, non-vocal learning animal. The calls’ rhythmic structure shows a progressive *rallentando* — a systematic slowing driven by changes in syllable and silence durations and the intervals between syllable onsets. The exact patterns of this *rallentando* are indicative of the caller’s sex and individually specific. These results reveal how seabird communication is shaped not only by efficiency universals, but also the specific pressures of colonial life. Our work highlights the temporal structure as an important axis of communication evolution, but also serves as a reminder to consider the species’ ecological reality and the function, not only presence, of temporal organisation.

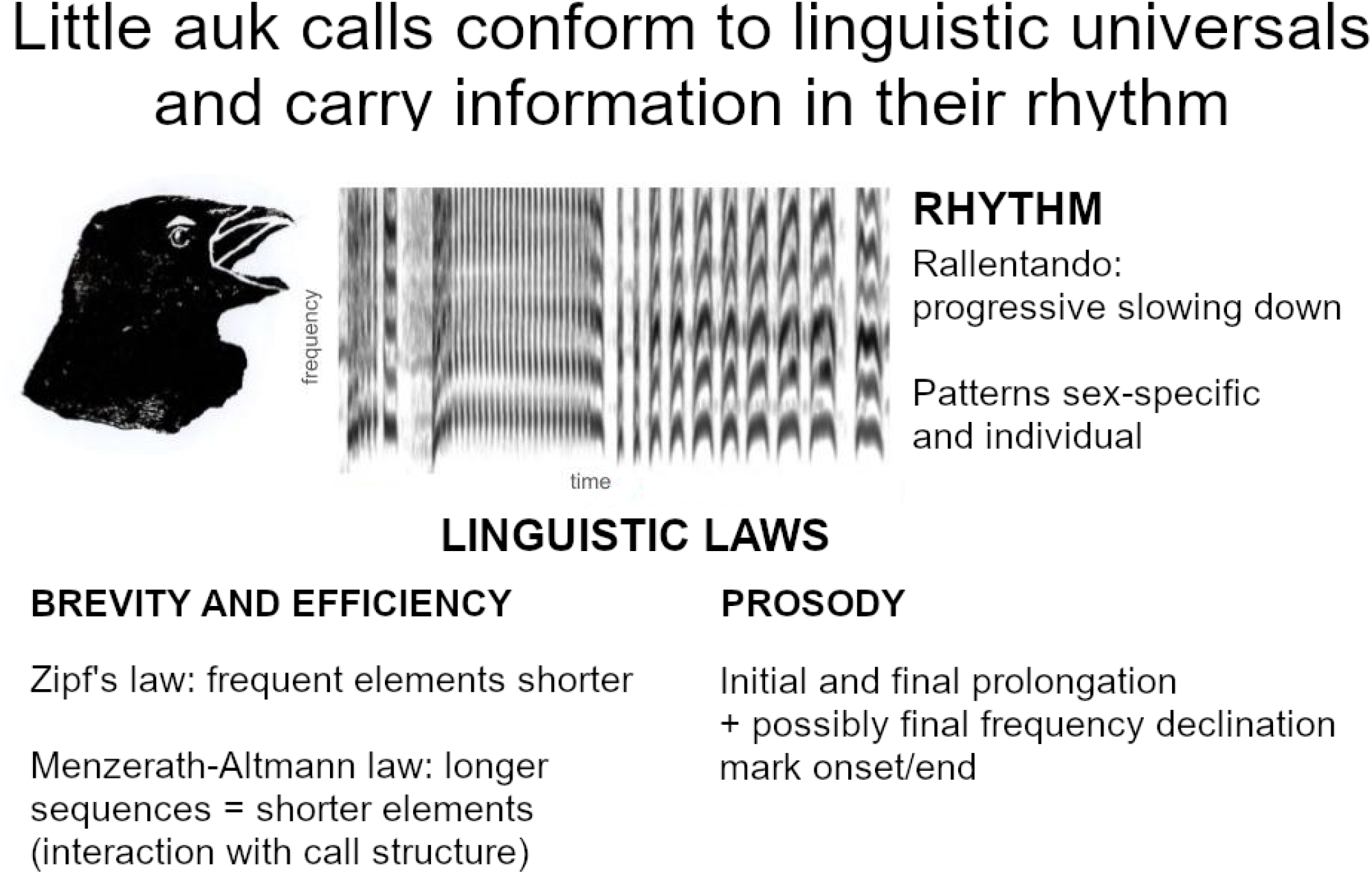

## Introduction

From birdsong to poetry, complex vocal signals are widespread across the animal kingdom. Sentences, songs, and composite calls can encode more information than simple one-syllable utterances [1], and repetition of an element bring more attention to it and increase the chances that it is in fact heard [2]. In order to be understood, complexity likely requires conformance to some organisational laws shared at least within the species.

Many human languages, in all their diversity, share not only grammatical universals [3], but certain statistical [4] [5] [6] [7] and prosodic [8] [9] [10] organisation. Likewise, human music shows reoccurring rhythmic patterns throughout cultures and traditions [11]. Might these human universals reflect a more general pattern in the evolution of communicative signals? In other words, are there any ground rules shaping the temporal organisation of communication across species?

Linguistic laws commonly refer to statistical patterns shared across multiple human languages. Over the last decades, they have attracted increased attention from biologists [7]. Some general patterns do in fact seem to emerge across not only communication strategies, but the organisation of life as a whole [7]. Linguistic laws addressing the coding efficiency of structure or information compression can describe many (but not all) communicative sequences produced by non-human animals. For example, Zipf’s law [4] predicts shorter forms for more frequent elements, and the Menzerath-Altmann law [5] predicts shorter constituent elements within larger constructs (Figure 1). Since vocal activity can be quite energetically expensive, these organisational rules minimise production costs for the speaker, caller or singer.

**Figure 1:**
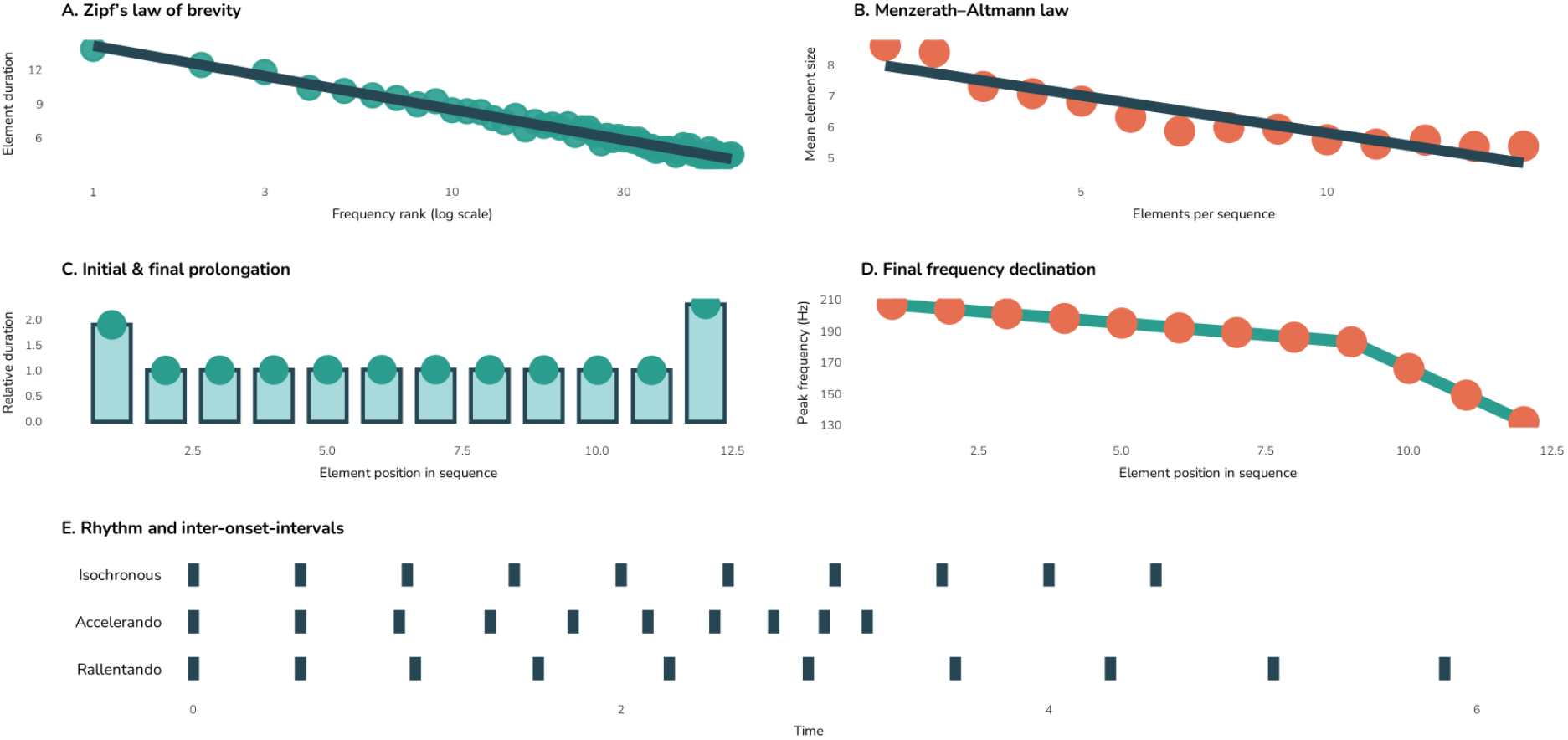
Visual representation of linguistic laws based on brevity (A-B) and prosody (C-D), as well as basic rhythmic organisation types (E) found in communicative sequences.

Communication, however, is structured by more than simple efficiency, and temporal markers can play crucial perceptual and functional roles. Prosody refers to intonation or rhythm beyond word meaning. These functions include clear demarcation of utterance boundaries. Final lengthening is present in many human languages [9] and some animal signals, such as the alarm calls of colobus monkeys (*Colobus polykomos* and *Colobus guereza*) [12] and budgerigars’ (*Melopsittacus undulatus*) speech-like utterances [13]. In the frequency dimension, such demarcation can present as final frequency declination [8] [14] [15]. This marks the end for listeners. Recent work highlights also an initial lengthening across human languages [10]. Together, these temporal variations segment signals and signal “when to listen”.

Another way to characterise temporal organisation is through rhythmic structure — the patterned relationships among element timing within a sequence [16] [17]. Rhythms can be described mathematically using rhythmic ratios [18], which capture how the timing of one element relates to another. Such analyses reveal whether a sequence exhibits isochrony (elements are spread in roughly equal intervals), tempo variation (such as *accelerando* or *rallentando*, where the sequence speeds up or slows down, respectively), or tem-poral irregularity (arrhythmicity; Fig. 1).

While both these approaches (linguistic laws and rhythm analysis) can be taken as simple descriptions of presence/absence in a species, recent evidence indicates that the temporal organisation of communication can interact with and be shaped by ecological, social, and sensory constraints [19] [20]. Isochronous rhythms have been suggested to carry information [21], and indeed various aspects of rhythmic organisation have recently been shown as information coding channels: isochronous patterns can carry meaning [21], and rhythmic cues can indicate caller identity [22, 23] or even sex [24]. Recent studies further suggest that the temporal organisation of vocal signals may be shaped by ecological, social, and sensory constraints [19] [20], underscoring the adaptive potential of timing itself. Thus, to understand the origins and meaning of temporally complex structures, we need to consider them at various levels, with integrative approaches combining linguistic, rhythmic, and biological perspectives.

The little auk (*Alle alle*) offers an ideal model for such inquiry. Despite the logistical challenges of studying this pelagic, High Arctic seabird [25], long-term research has produced extensive multidimensional datasets on its ecology, social behaviour, and communication [25]. Little auks lead social lives in dense breeding colonies [26], which is expected to lead to complex and efficient communication systems [27] [28]. Little auks rely heavily on vocal communication [29] and exhibit a complex repertoire [30], with calls carrying cues to e.g. identity [22], partnership [31], and emotional status [30] [32] of the caller, yet spectral analyses failed to show sex-specific patterns in their vocalisations [31]. The species uses a single complex call type - the *classic call* [30] - is composed of three syllable types (in rare cases four; Figure 2), and shown to carry multiple information types [30] [22] [31] over large distances [33]. While their context-specific functions are not yet fully understood, it is recognised as a socially crucial call. Importantly, seabirds are not considered vocal learners, thus avoiding any human-animal mimicry loops [34] in their vocalisation patterns.

**Figure 2:**
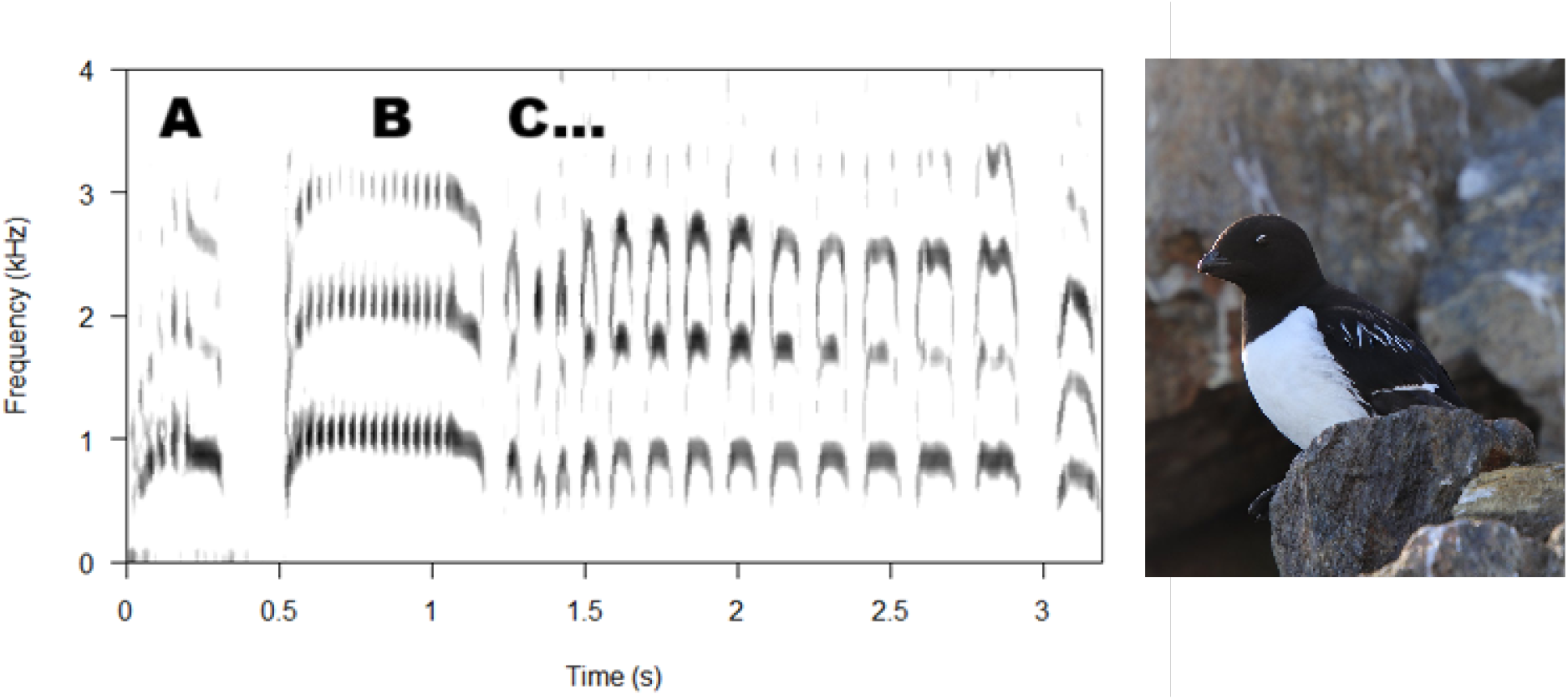
Little auks’s *classic calls* dominate the summer soundscape of the High Arctic. The call is comprised of three syllable types (A, B, and a series of brief C syllables), but in rare cases an additional, final syllable (D, similar in structure to A) is used. Photo credit: Marion Devogel

Here, we examine the temporal organisation of the little auk’s *classic call* in detail. Based on the syllable durations, inter-onset-intervals and silences between each two consecutive syllables, we test whether these calls conform to established linguistic laws observed in human communication, and whether their rhythmic structure reveals consistent patterns that may encode biologically meaningful information.

## Results

### Menzerath-Altmann law

After controlling for the individual ID, but not syllable type, the calls conformed to the Menzerath-Altmann law (larger constructs were composed of shorter syllables; Fig.3, panel A). When considering each syllable type separately, this did not hold: only syllable C showed a slight, but not significant, decrease with the call’s size (Figure 3, panel D).

**Figure 3:**
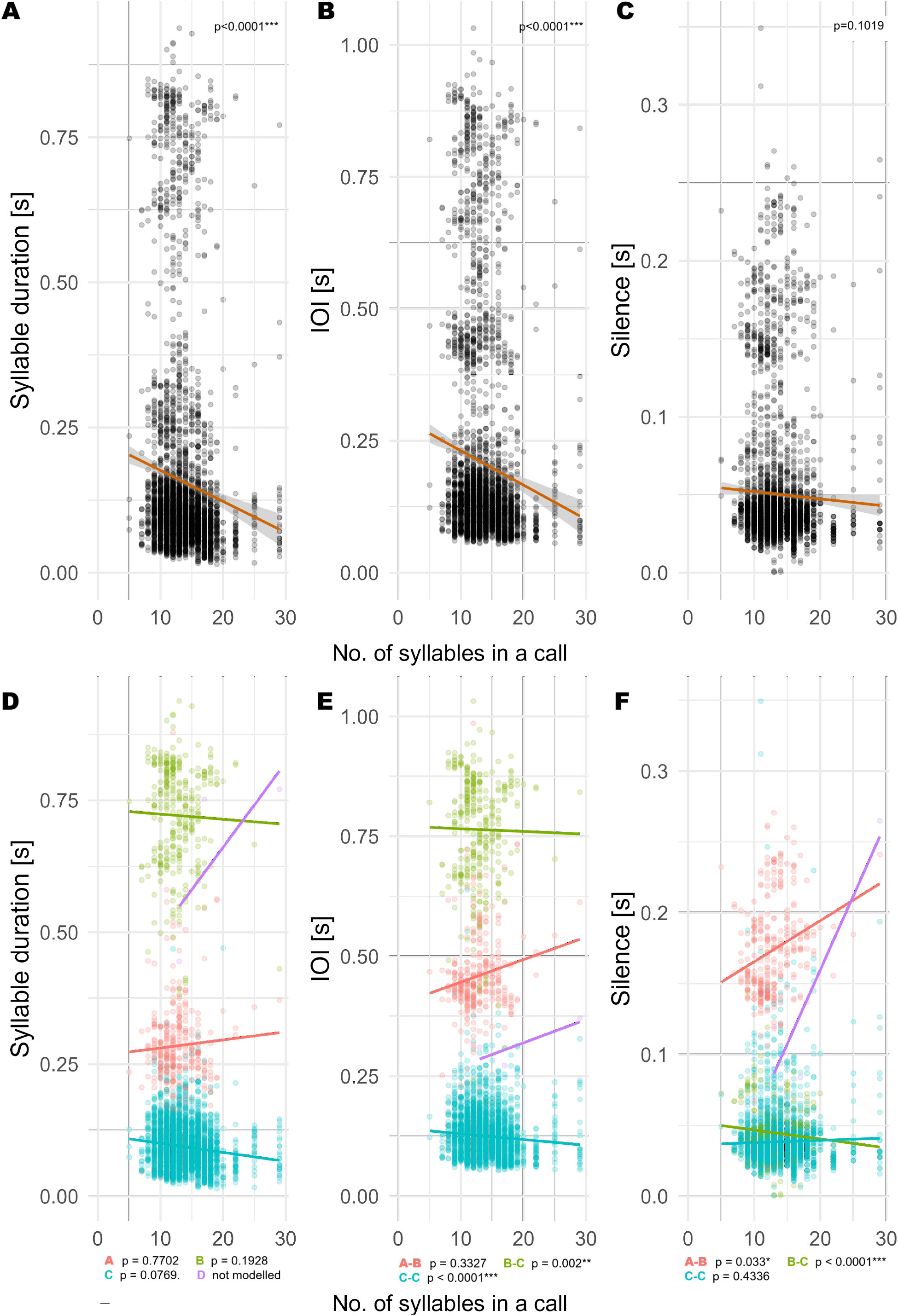
Linear models were used to test conformance of the classic call to Menzerath-Altmann law. Panels A-C consider all elements in a sequence, and D-F investigate whether patterns hold within syllable types. Syllables D are very rare and thus not modelled.

The inter-onset-intervals (IOI) conformed to the Menzerath-Altmann law both within the entire construct (without controlling for syllable pair type) and for intervals connecting syllables B-C and C-C (Figure 3, panels B and E). Silences between the syllables were not overall shorter in larger constructs (Figure 3, panel C), but did significantly decrease for B-C pairs and increase for A-B pairs (Figure 3, panel F).

### Initial and final lengthening

After correcting for the callers’ identity, little auk *classic calls* conformed to both initial and final lengthening laws in all measured aspects: syllable duration, IOI, and silence (Figure 4; p values for all measures ≤ 0.0001).

**Figure 4:**
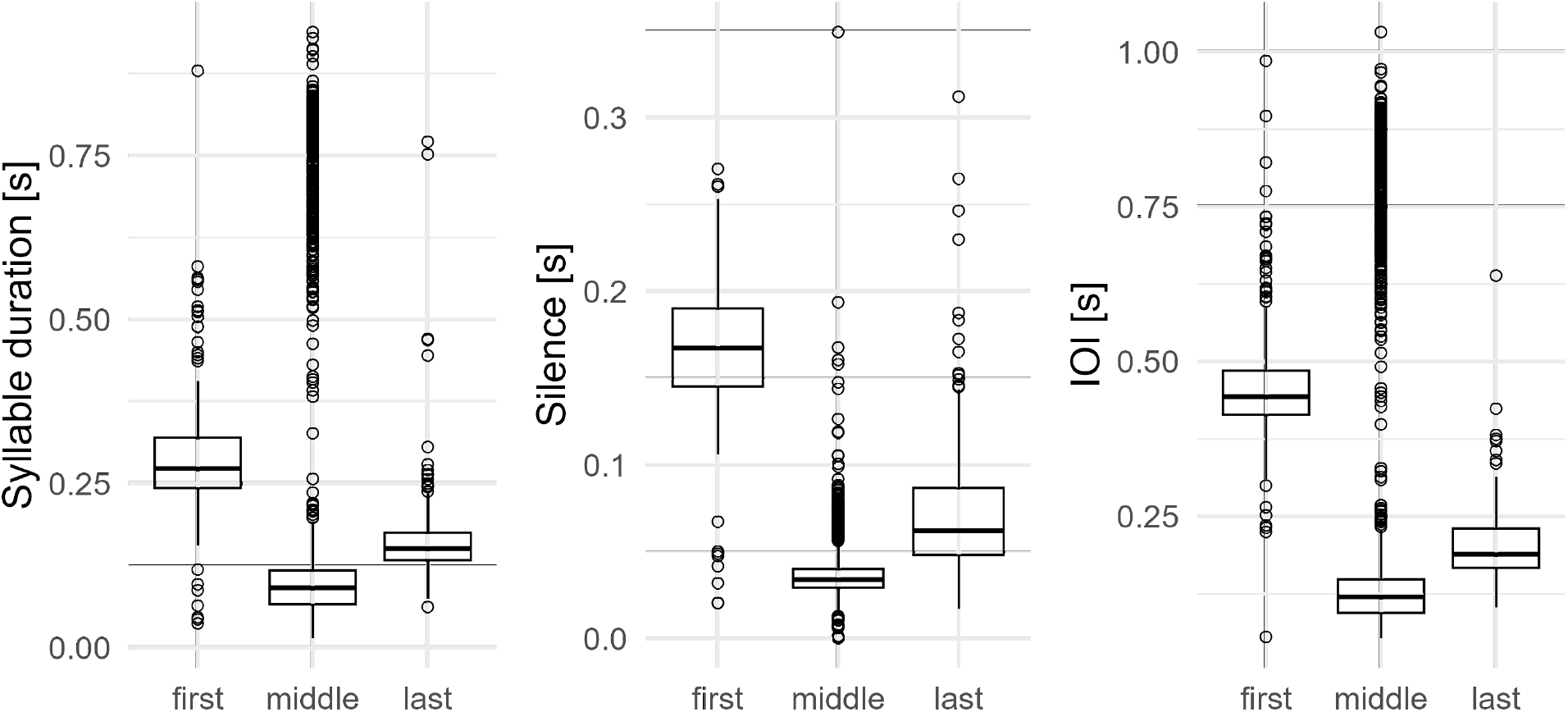
Linear mixed models’ results at the scale of the entire sequence indicate initial and final prolongation in all measured parameters (syllable duration, IOI, silence).

### Rhythmic ratios

To interpret the integer ratios (rk values), it should be mentioned that values of 0.5 indicate that the two compared elements are similar, values higher than 0.5 indicate that the lengths of the consecutive elements decreases as the sequence progresses (*accelerando*: the syllable, IOI or silence is shorter than the previous one), and values below 0.5 indicate the opposite (*rallentando*: the syllable, IOI or silence is longer than the previous one).

All local and global measures indicated a *rallentando* in the call (Fig. 5). The local rhythmic ratios based on silence durations were particularly low, indicating that inter-syllable silences lengthen most as the sequence slows down (Fig.5). On a global scale, however, the silence-based rk values seem to become more balanced (Fig.5).

**Figure 5:**
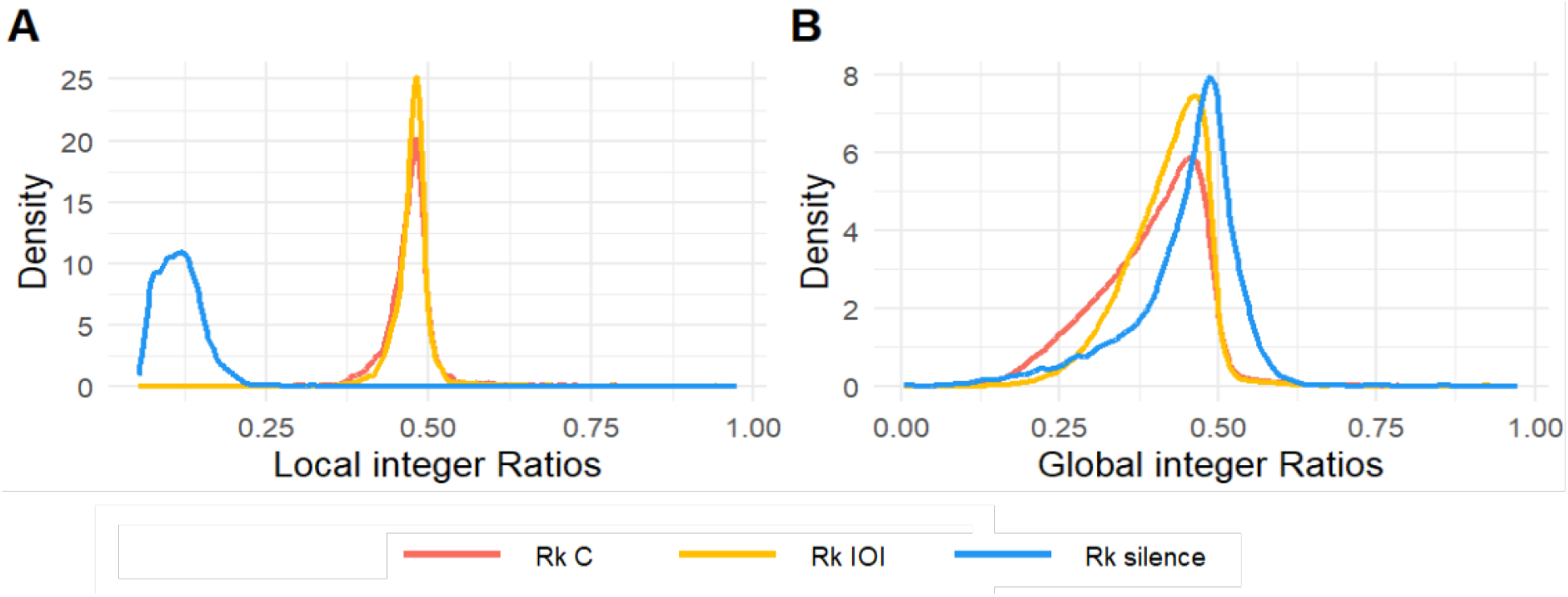
Densities of local (A) an global (B) integer ratios based on syllable durations (red), inter-onset-interrvals (yellow) and silences (blue).

Different organisational patterns emerged when the syllable order (for local values) and relative distance between the syllables (for global values) were considered (Figure 6). Local rhythmic ratios based on syllable duration and silence increased, and these based on IOI decreased as the call progressed, although all values stayed in the rallentando range (regression line below 0.5, Figure 6, panels A-C). All global rhythmic ratios indicated a clear, progressive rallentando (values decreased as the sequence progressed; Figure 6, panels D-F).

**Figure 6:**
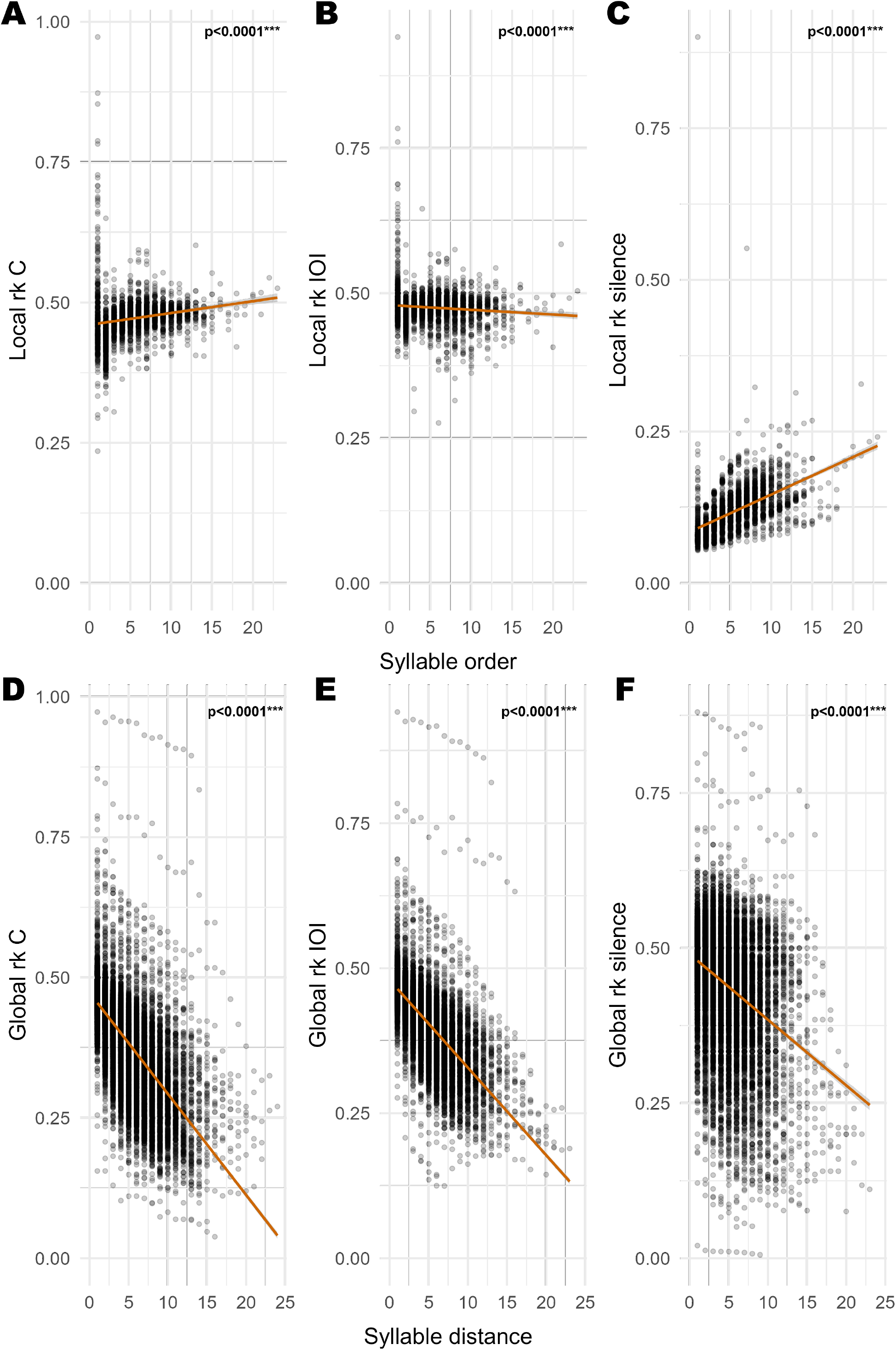
Rhythmic structure of the C-syllable part of the *classic call* shown as local (A-C) and global (D-F) integer ratios. Global ratios show a clear, progressive *rallentando*.

### Information coding in rhythmic structures

Using permuted discriminant function analysis (pDFA, Figure 7, panel B), vocalisations could be reliably assigned to sex (Figure 7, panels A-B) and individual (Figure 8, panels A-C, Figure 7, panel B), but not nesting pair, based on the local rk values (Figure 7, panel B).

**Figure 7:**
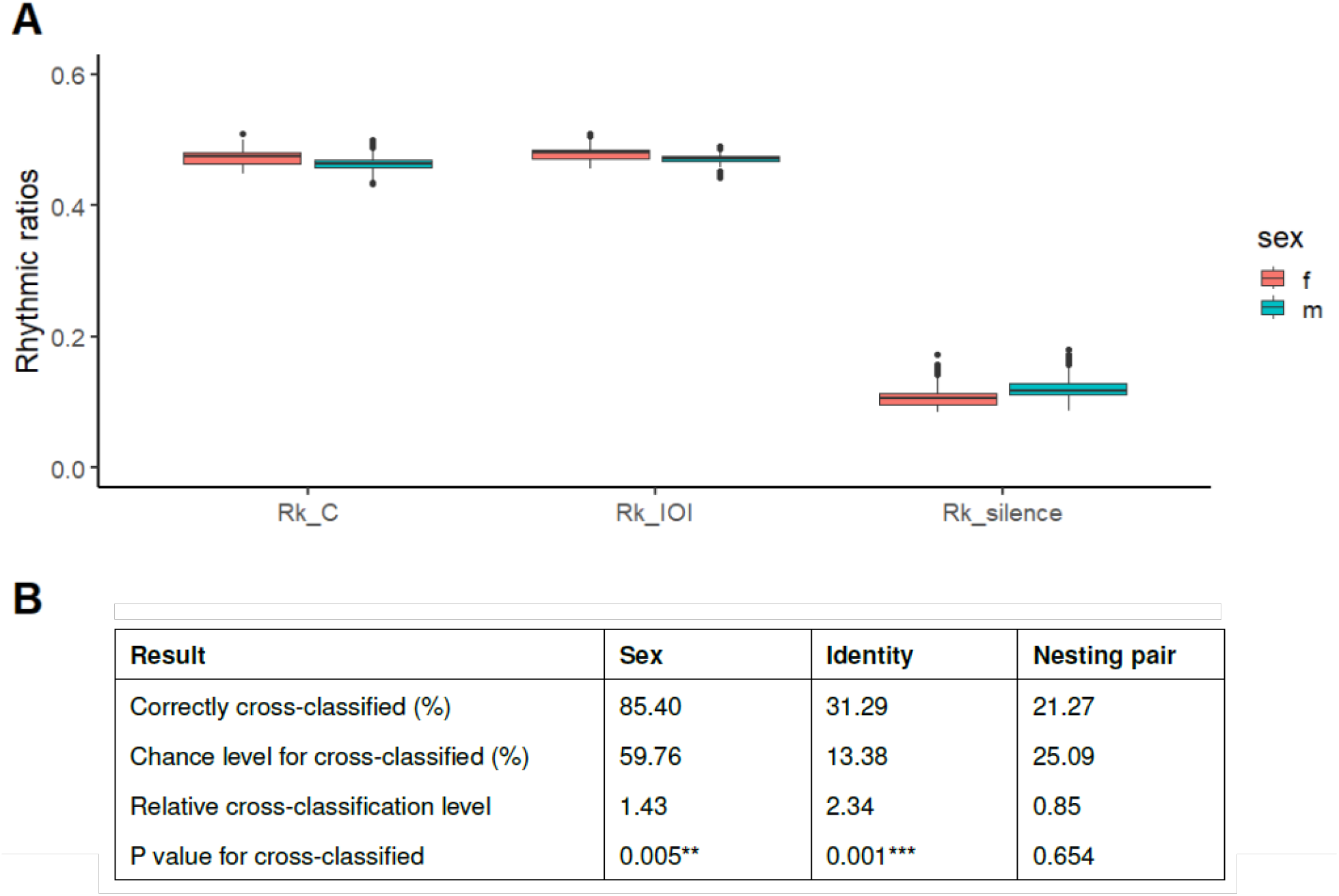
Local rhythmic ratios differ between sexes (A) and based on their values calls can be correctly assigned to both sex and caller, but not nesting pair (B).

**Figure 8:**
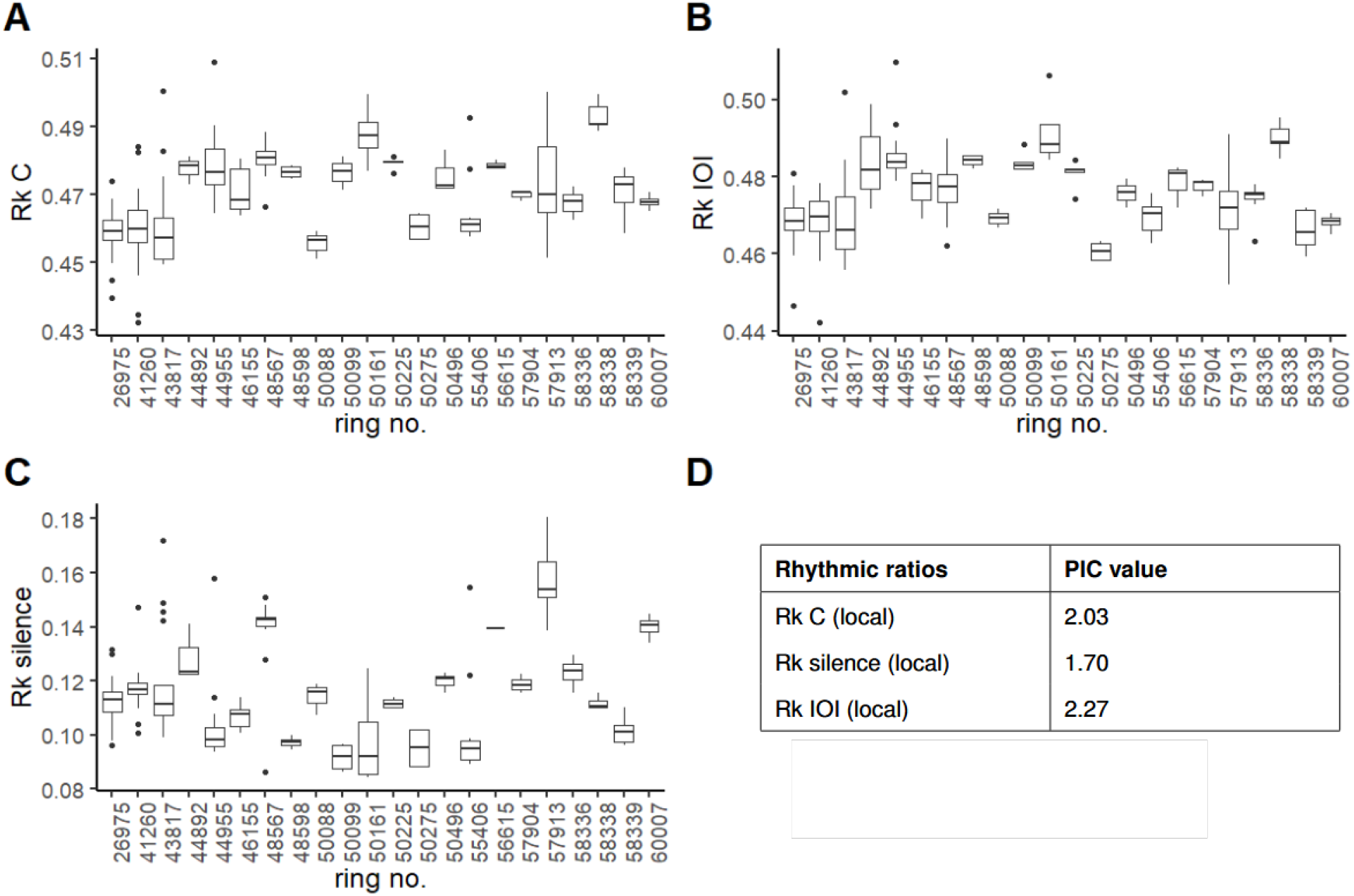
Individual distribution of local rhythmic ratios (A-C) and their potential for identity coding (D).

Potential of identity coding (PIC) values over 1 indicate importance for identity coding, and values over 2 that a parameter is very important. Each of the local rhythmic ratios showed importance for identity coding (Figure 8, panel D). Individual variation within each of these parameters seems to also play a role (Figure 8, panels A-C).

## Discussion

The little auk is the first non-human, non-vocal-learning species in which conformance to multiple linguistic laws, including both simple vocal efficiency and prosodic patterns, has been demonstrated. Their *classic calls* show a *rallentando* rhythmic structure, and rhythmic differences are both individual and indicative of the caller’s sex, previously not identified acoustically with success in the species.

### Vocal efficiency and prosody

Little auks’ *classic calls* conform to the tested linguistic laws, including both vocal efficiency (Menzerath-Altmann law) and prosody (both initial and final lengthening across syllables, inter-onset intervals, and silences).

We explicitly excluded two other linguistic laws from formal testing. First, Zipf’s law of abbreviation predicts shorter forms for more frequent elements [4]; this did not require separate analysis because the classic call’s structure already embodies it—the brief C syllables are the only repeated elements and thus most common.

Second, we omitted fundamental frequency declination analysis, as it concerns spectral rather than temporal structure. Fundamental frequency (f0) declination (decrease) is observed in statement-type utterances of human languages [8] as well as certain non-human primates (vervet monkeys *Cercopithecus aethiops* [14], rhesus macaques *Macaca mulatta* [14], baboons *Papio cynocephalus ursinus* [15]), and budgerigars [13]. While not explicitly tested, terminal C syllables do show lower fundamental frequency in comparison to other elements in the sequence, accentuating the call’s end and likely signalling the end of a sequence [35]. This shows a further accentuated end to the utterance rather than an overall decrease throughout the vocalisation, similarly to the pattern observed in budgerigar song [13] and unlike the gradual primate f0 declination patterns [35]. Given the vocal complexity of avian systems and the vast existing recording libraries for this group, we recommend revisiting this question from a more evolutionary perspective, comparing the final element’s frequency patterns across species and social systems.

Most studies considering linguistic laws in non-human animals concern laws of brevity (various Zipf’s laws, Menzerath-Altmann law), which translate to vocal efficiency, i.e. optimising the energy expenditure and information compression. Conformance to the laws of brevity is unsurprising in nature - they describe sequences from genes and proteins to sentences and books [7]. Prosody is somewhat more “sophisticated”, engaging cognitive processes and signaling the start and end of a sequence, as well as contributing to meaning and emotional expression in humans [36]. Interest in non-human prosody is rarer, but growing [35], and birds have been shown to perceive and react to prosody and grammar modifications [37] [38]. Interestingly, while vocal efficiency in little auk calls showed varying results depending on whether the full call or parts of it were considered, the prosodic initial and final lengthening were consistently found across syllables, intervals, and silences. So far, conformance to both f0 declination and final lengthening has only been described in humans and budgerigars [13]. To our knowledge, little auks are the first non-human, non-vocal-learning species for whom conformance to multiple linguistic laws has been demonstrated, including both vocal efficiency and multiple prosodic patterns. This indicates a complex signaling system in a non vocal learner species evolutionary and behaviourally far from humans.

Does all this mean that little auks use “language”? Most probably not. Even though they are not universal, compression and efficiency are general rules in animal signals [39]. While conformance to basic vocal efficiency laws is still often called “language-like”, such organisational rules can arise without the rich combinatorial meaning and flexibility that characterise human language, or communicative sequences of some other species [40] (most notably other primates [41] and vocally learning birds [42]). What is interesting, is that the Menzerath-Altmann law describes both full sequences and certain parameters within the calls (IOI of B-C and C-C pairs, as well as silences between B-C pairs; Figure 3). Within the syllable pairs, we argue that the IOI shortening in longer sequences in fact represents the efficiency rule (sequences composed of more elements are characterised by shorter elements - in this case, the inter-onset-intervals). Both syllable durations within syllable types, and the variation in syllable C duration are highly individual [22] and might not be up to efficiency adjustments - it is hence understandable that they remain unaffected by sequence length. At the same time, the silence between syllable A and B increases, and silence between syllable B and the first C syllable decreases with sequence length across individuals. Breathing patterns and vocal output are closely related and mutually influential [43]. While prolonged silence between the first and sec-ond (A and B) syllable could be a result of a breathing limitation (“a longer pause is needed between a long syllable and the following series”), the roughly 0.12-0.25 s range of these silences might not be sufficient for a deep breath, and thus possibly relate to the prosodic organisation.

Being more perceptual, the case of prosody is particularly interesting. When listening to non-word, non-sensical utterances, human infants react to correct prosodic patterns, indicating an early attentiveness to elements signaling a communicative sequence [44]. This might suggest that in species with a prominent prosodic organisation of communicative signals, it is expected already from very young listeners without extensive experience. Little auks might represent a similar care: while seabirds are not vocal learners, they live extremely socially complex lives, often guided by intense vocal interactions (e.g. [30] [45] [46]. In a crowded colony of very vocally active birds, prosody might prove a necessary guiding tool to the listener, indicating where information-heavy sequences might begin and end. While it remains unclear how prosodic patterns might be perceived and used in this group, future playback experiments might shed some light on the question. Circling back to the paragraph’s initial question: out results stress that vocal communication in general tends to follow more (although not completely) universal patterns, and that complex organisation similar to some human languages can be found in poorly considered systems, such as seabirds.

The fact that a non-vocal-learning seabird shares multiple organisational rules with human languages raises some important evolutionary questions: did these patterns emerge analogously under similar pressures, were they inherited from a shared ancestor and partially lost in some lineages, or did they influence each other? Humans are one of the few mammal species that vocally learn, and are in fact expert vocal learners [47]. Unlike many species, we exhibit cortical sensitivity to animal vocalisations in general, not just conspecific ones [48], and mimicking animal sounds has been proposed as one contributor to the emergence of human language [49]. From function to production mechanisms and neural pathways, we share a remarkable amount with vocally learning birds [50], and Tsing [34] highlights how mutual human-animal mimicry can create feedback loops where both sides shape each other’s behaviour. Given that little auks are extremely unlikely to have been influenced by human speech, however, any resemblance to human prosodic organisation must arise from shared functional pressures or conserved ancestral features.

The discovery that a non-vocal-learning seabird exhibits both prosodic timing and linguistic efficiency laws extends the search for communication universals beyond vocal learners and primates [50] [7]. Emotional and interactional prosody—similar timing patterns coordinating social exchanges—has been proposed as a precursor to linguistic prosody across mammals, songbirds, and humans [51]. Our results demonstrate that such temporal organisation can emerge in innate vocal systems, suggesting these patterns arise from general communicative pressures rather than vocal learning or specialised language faculties. Human prosody may thus represent one example of broader solutions to the challenges of efficient, intelligible signalling [50].

### Rhythmic organisation and information

In the little auk, vocal exchanges between partners mediate the duration of incubation duties [29], and likely play a large role in social organisation via individual [22] and group [31] signatures. The species does not exhibit a clear sexual dimorphism, and previous work indicated a lack of sex signatures in the spectral content of their calls [31]. Our results therefore reveal a previously overlooked channel: information on the caller’s sex is encoded not in the spectral structure, but in the rhythmic organisation of the *classic call*. Since this information is not present in other dimensions of sound, it does not seem to be redundancy-driven. Male vocalisations slow down at faster pace than female ones, but this is less silence-driven than in females (that is: syllable durations and IOIs extend more and silences extend less in males than females as the sequence progresses). If the birds use these differences to aid them in identifying a potential mating partner, this differential was likely driven by sexual selection over time.

Rhythmic organisation is increasingly recognised as a fundamental feature of animal vocalisations (e.g. [52] [53], [19]). Recently, it has been shown that rhythmic patterns can carry information on e.g. caller’s identity, sex or population of origin within isochronous [23] [54] and arrhythmic [19] sequences. The accelerating ecstatic displays of the African penguin (*Spheniscus demersus*) have been suggested to play a role in quality or motivation signaling [55]. In contrast, the little auk *classic calls* slow down, and this *rallentando* pattern is characterised by a stable, linear decrease in the global rk values. We have only considered local values as information-carrying channels, since they do not follow an obvious linear decrease. Indeed, all local rk parameters (whether based on syllable duration, IOI, and silence) achieved high potential of identity coding (PIC) values, and permuted discriminant function analysis reliably classified both sex and individuals based on rhythmic parameters alone. We observed that the extent to which rhythmic ratios vary for each individual varies substantially between individuals. This supports the growing body of evidence that the extent of variation is in itself a very individual trait [23] [19] [22].

This complex temporal organisation adds another information dimension to the *classic call*, which emerges as the species’ most reliable channel for advertising socially important information on the individual [31] [22], possibly transmitted at large distances [33]. Colonial seabirds face extreme acoustic interference from dense breeding aggregations, where overlapping calls saturate the soundscape. The *classic calls* used in these study were recorded from stationary birds, but they are also produced in flight [30], where their frequency content are likely distorted by the Doppler effect [56] [57]. While Doppler shifts themselves have been suggested to play a role as proximity cues in seabirds (blue petrels *Halobaena caerulea* and Antarctic prions *Pachyptila desolata*) [58]), we suggest that the temporal or rhythmic dimension could also help maintain the informational content of a call despite frequency distortions. Temporally structured calls with clear boundaries and informative rhythms thus likely enhance social recognition and coordination, while extended terminal silences provide perceptual space amid potentially overlapping neighbours’ calls.

### Conclusions

We provide the first evidence that seabird communication shows a complex temporal structure, with similarities to human communication going beyond vocal efficiency. We also highlight the importance of rhythmic organisation in information coding in seabirds. These results stress the complexity of seabirds communica-tion, shaped by the pressures of colonial life.

## Methods

### Ethical note

This study was based on readily available passive acoustic recordings collected by the Polar Ecology Group, University of Gdańsk, over 2019-2023, and as such did not require a specific permit from an animal research authority. The annual fieldwork, performed in the little auk breeding colony located at the aread of National Park (Hornsund, Spitsbergen), was conducted under the lead of KWJ, with annually granted permissions of the Governor of Svalbard, and following all relevant regulations and best practices in animal research [59]. Each season, both individuals from each nesting pair recorded in the study were sexed, ringed and recorded inside the nests.

### Data availability

All acoustic data, except for the 2023 set, have previously been published in [22]. Full data and scripts generated in this work, as well as supplementary materials are available at: https://osf.io/3wuxb/?view_only=c6fbbb118cec4e07a21fbc450eef4713

### Data preprocessing and calculation of indices

All recordings have been manually screened by ANO and GV in Raven Pro to extract good quality *classic calls* and assign them to known (sexed, and ringed) individuals based on the video footage taken at the entrance to each nesting cave (for details consult [22]). Each call was then manually annotated in Raven Pro by ANO, to mark the start and end times of each syllable, as well as syllable type.

The obtained selection tables were then used with a custom-written R script published with the project data, based on [22] and [16], to calculate the duration of each syllable and its location within the sequence, as well as six different rhythmic ratios (rk) (Table 1). Since the first part of the call (A and B syllables) is clearly different from the C-syllable sequence, rhythmic ratios (rk) were calculated for the C-syllable durations, their inter-onset intervals (IOIs) and silences only.

**Table 1:** Definitions of the temporal parameters calculated for each vocalisation.

| Abbreviation | Definition |
| --- | --- |
| Rk IOI (local) | Integer ratios based on inter-onset-intervals between C syllables, calculated as a ratio between the IOI's duration and the sum of this and the following IOI's duration [18]. Note that the same calculation is sometimes described as "mean acceleration rate" [55] $r_k = \frac{\text{ioi}_{-1}}{\text{ioi}_{-1} + \text{ioi}_1} \quad (1)$ |
| Rk C (local) | Integer ratios based on C syllable duration, calculated as a ratio between the syllable's duration and the sum of this and the following syllable's duration |
| Rk silence (local) | Integer ratios based on inter-syllable silences, calculated as a ratio between the silence's duration and the sum of this and the following silence's duration |
| Rk IOI (global) | Global integer ratios [16] based on inter-onset-intervals between C syllables, calculated not only for adjacent intervals in the sequences, but for all interval pairs of a sequence $r_k^{(n,m)} = \frac{\text{ioi}_n}{\text{ioi}_n + \text{ioi}_m}, \quad \text{for all } n, m \in \{1, \dots, N\}, n \neq m \quad (2)$ |
| Rk C (global) | Global integer ratios [16] based on C syllable duration |
| Rk silence (global) | Global integer ratios [16] based on inter-syllable silences |

### Data analysis

#### Menzerath-Altmann law

To check whether an increase in size of a vocalisation (number of syllables within it) leads to a decrease of its elements’ size, we ran three sets of linear mixed models, based on (1) syllable duration, (2) IOI duration, and (3) silence duration as elements (response variable). Each set included (1) a full model including all syllable types or syllable pairs (metrics calculated for two consecutive syllables), and (2) three models for a single syllable or syllable pair type. The models used the function *lmer* of *lme4* package ([60]) with a *bobyqa* optimizer (fitted using *allFit* function of *lme4* package), with the sequence length (number of elements) as a fixed factor, and the individual ring numbers (birds’ identity) as a random effect grouping factor. Sex was not considered, since it resulted in errors caused by model overfitting. Syllables D and syllable pairs including D syllables were not modelled, since too few datapoints were available. To calculate the p-values, each model was accompanied by a small model excluding the sequence length; the models were then compared using the *PBmodcomp* function of *pbkrtest* package ([61]).

#### Initial and final lengthening

To check whether little auk *classic calls* follow the initial and final lengthening laws, we ran a set of thee linear mixed effects models based on (1) syllable duration, (2) IOI, and (3) silence. The models used function *lme* of *nlme* package) with the element location (first, middle or last) as a fixed factor, and the individual ring numbers (birds’ identity) as a random effect, estimating a separate residual variance for each position group as a heteroscedasticity fix (data first inspected with a Bartlett’s test, using *bartlett*.*test* function of the *stats* package). We then ran pairwise comparisons using the *emmeans* function of *emmeans* package ([62]). This was considered at the scale of the entire sequence, not controlling for syllable type.

#### Rhythmic patterns

To look for patterns in the distribution of rhythmic ratios in the *classic call*, we ran a set of six linear mixed models based on the available rk values (Table 1), with the rk values as response factors, position in a sequence as a fixed factor, and the individual ring numbers and sex as random effect grouping factors. All models used the function *lmer* of *lme4* package ([60]). Each model was accompanied by a small model excluding the position in a sequence; the models were then compared using the *PBmodcomp* function of *pbkrtest* package ([61]).

#### Information coding in rhythmic patterns

To see whether the rhythmic patterns of the *classic call* carry information on the caller’s (1) sex or (2) identity, and whether (3) nesting partners use similar rhythms, we ran three permuted discriminant function analyses (pDFA, [63]) models in nested design: (1) sex as test, ring number as control; (2) ring number as test, sex as restriction; and (3) nest as test, ring number as control. Each model was ran with 1000 permutations, using mean local rk values for each vocalisation and custom functions by Mundry, based on the *MASS* package ([64]).

To investigate the importance of each rk type for vocal individuality, we calculated the potential of identity coding (PIC, [65]) of the mean local rk values, using the *calcPIC* function of *IDmeasurer* package ([66]).

## Authors contributions

ANO: Conceptualization, methodology, formal analysis, investigation, data curation, writing: original draft, review and editing, software, visualisation, project administration, funding. KWJ: Conceptualisation, investigation, writing: review and editing, funding. LSB: Methodology, software, writing: review and editing, supervision.

## Acknowledgements

Heartfelt thanks to Guillaume Verchère, who processed a substantial part of the recordings; everyone who contributed to the fieldwork and data processing over the years; the expedition members of Polish Polar Station Hornsund for their ongoing support; Feliksa Żurawska and Blank Kwiatkowska for little auk art; and the little auks without whom this work would not be possible.

